# Polyandry is extremely rare in the firefly squid, *Watasenia scintillans*

**DOI:** 10.1101/2019.12.13.875062

**Authors:** Noriyosi Sato, Sei-Ichiro Tsuda, Nur E Alam, Tomohiro Sasanami, Yoko Iwata, Satoshi Kusama, Osamu Inamura, Masa-aki Yoshida, Noritaka Hirohashi

**Affiliations:** Oki Marine Biological Station, Shimane University, 194 Kamo, Okinoshima, Oki, Shimane 685-0024, Japan; Department of Fisheries, School of Marine Science and Technology, Tokai University, Shizuoka 424-8610, Japan; Department of Applied Life Sciences, Faculty of Agriculture, Shizuoka University, 836 Ohya, Shizuoka, Shizuoka 422-8529, Japan; Atmosphere and Ocean Research Institute, The University of Tokyo, 5-1-5 Kashiwanoha, Kashiwa, Chiba 277-8564, Japan; Uozu aquarium, 1390 Sanga, Uozu, Toyama 937-0857, Japan

**Keywords:** monogamy, monoandry, cephalopods, squids, deep-sea, spermatangium

## Abstract

Although polygamy has versatile benefits for both sexes, many species favor monogamy for reasons with the clarity or unclarity. In cephalopods, all species are regarded to be polygamous, which could be attributed to their common life-history traits. Contrary to this prediction, we show empirical evidence for monogamy in the firefly squid, *Watasenia scintillans*. The peak spawning season comes after male disappearance owning to long-reserved spermatangia deposited by male at exact locations (bilateral pouches under neck collar) on female with a symmetric distribution. Such a non-random placement of spermatangia prompted us to hypothesize that females engage in lifetime monoandry. Hence we assigned genotypes of female-stored spermatangia and offspring. We found that in 94.7 % females, the spermatangia were delivered from a single male and all embryos in the same egg string sired by sperm from stored spermatangia. Throughout the season, relative testes mass was much smaller in *W. scintillans* than all other cephalopods previously examined. The mean number of male-stored spermatophores was approximately 30, the equivalent to 2.5 mates. Our demographic and morphometrical data agree with the prediction that monogyny is favored when potential mates are scarce such as absence of female remating. Together, these results suggest the likelihood of mutual monogamy.

## 1. Introduction

In recent decades, research on behavioral and molecular ecology identified polyandry (a female mate with multiple males), a prerequisite for postcopulatory sexual selection, as a prevalent mating pattern across animal taxa [1-5]. In this pattern, females are able to receive direct or indirect benefits to a greater extent by mating with multiple males than with a male, resulting in offspring with ‘good genes’ or higher genetic diversity [6, 7]. Conversely, costs of polyandry (or polygamy) are borne on both sexes such as increased risks of predation, disease/virus infection, harassment during courtship or copulation, and therefore often shortening their lifespan [8-10]. Yet, cost-benefit trade-offs in choosing a mating pattern cannot be easily anticipated due to the underlying social and population complexities combined with various life-history and reproductive strategies. Furthermore, even though polygamy bears high costs on each sex, it would not necessarily be transitioned to the contrasting mating pattern, monogamy (both male and female mate with one partner). Because monogamy, as a whole, should meet the criteria of either ‘mutual benefits’ for both sexes [7, 11, 12] or ‘unilateral benefit’ to one sex over the other (e.g., postcopulatory mate guarding). The latter could also generate sexual conflict or energetic costs on the guarders [13, 14]. Hence, monogamy is favored only in environmental conditions where the opportunity or benefit to monopolize mates does not exist [15]. For examples, birds and mammals have entailed biparental care for young and long-term (prolonged) pairing, setting a precondition for evolution of monogamy [14, 16, 17]. Species that are under constraints of feeding or breeding habitat often choose the monogamous relationship to protect these resources [18, 19]. Female disperse could be a reason for monogamy because searching for extra-pair mates is costly and risky for males [20]. However, monogamy could have arisen in species without these cases, therefore still posing an evolutionary puzzle [21].

Most cephalopod species have a short lifespan (approximately one year), reproduce semelparously and display a diverse array of mating behaviors in favor of adaptive or alternative consequences of intrasexual reproductive competition [22-25]. Behavioral and anatomical observations of some coastal species have uncovered the coherent reproductive tactics and traits thereof in the context of promiscuous mating [22, 26-29]. Notably, all cephalopod species reported are regarded to be polyandrous [30-33], except that the diamond squid, *Thysanoteuthis rhombus* appears to form the long-term pair-bond during migration and are therefore possibly monogamous [34]. Although this phenomenon needs confirmation with genetic analysis, many other species also require further investigations to determine whether the influence of observed female promiscuity is limited to their mating behavior, or even reached to their offspring paternity. Since it is now widely recognized that apparent mating behavior, regardless of whether it is promiscuous or monogamous, does not always coincident with, or proportional to, genetic parentage [3, 35].

In cephalopods, the main factors that could possibly influence to this inconsistency are cryptic female choice (CFC), last-male sperm precedence (LMSP) and extra-pair copulation as a consequence of alternative reproductive tactic (ART) [24, 25, 33]. CFC is the postcopulatory mechanism by which females bias storage or usage of spermatozoa [36]. In the Japanese pygmy squid *Idiosepius paradoxus*, females strip off the spermatangia handed on from unfavorable males [37]. In addition, as females have smaller sperm-storing capacity (within the seminal receptacle) relative to sperm number in the spermatangia and sperm use from the seminal receptacle is under the female control, it is possible that CFC could also operate during these processes [38]. LMSP would be in practice if most recent mate takes a disproportionate share in the paternity [39]. In the cuttlefish, *Sepia lycidas* [40] and perhaps the dumpling squid, *Euprymna tasmanica* [41], males replace previously stored rival ejaculates for their own. Such the sperm displacement as well as widely observed mate guarding behavior must have evolved through LMSP [39]. ART refers to behavioral traits selected to maximize fitness in two or more alternative ways in the context of intraspecific and intrasexual reproductive contest [42]. Several loliginidae species show ART with distinct male morphs in mating posture and ejaculate traits [24, 43, 44]. Males with larger body mass (‘consorts’) deposit the spermatophores near the oviduct thereby sperm gain higher accessibility to eggs, whereas smaller males (‘sneakers’) take an alternate route to fertilize eggs by depositing sperm at either distal or proximal time from egg spawning [28, 45, 46].

Regardless of the presence of these male- or female-driven strategies that can facilitate post-mating paternity skew, it is uncommon, but not impossible, that one male can monopolize exclusive paternity of a brood if polyandry underlies the mating practices in a given species. In general, most cephalopods may have developed life-histories and reproductive modes suitable for pursuing polyandry; they are short lived, semelparous and capable of long-term storing multiple sperm packages in the female [33]. So far there has been no report, to our knowledge, for paternal care or provisioning by males after mating. Postcopulatory mate guarding is a possible cause of mutual monogamy, if it can deprive from both sexes the opportunities for seeking additional mates. Mate guarding by males is often seen in squids [44, 46, 47], however it is either temporal or incomplete, and sometimes interrupted by extra-pair copulations. Some organisms living under photic zone may suffer from low mate availability, another possible cause of monogamy, due to low population density [48], however mating pattern of any deep-sea species yet to be determined. Although there are many untested scenarios or conditions where monogamy is preferably selected, in the current consensus view, monogamy is seemingly unlikely to be a prevalent strategy in cephalopods [31, 32].

To date, however, most knowledge of reproductive ecology in cephalopods has gained from studies with limited representatives, mainly those with coastal habitats, because behavioral observations are possible in field or aquarium. Besides these approaches, the DNA finger printing has been a promising technique to track down the outcomes of mating events. To gain global insights into cephalopod reproductive systems, in this study, we chose the firefly squid, *Watasenia scintillans* because their ecological characteristics are rather clear [49], yet their reproductive mode remains largely equivocal. Furthermore, they are commercially important resources, which allows the fishery to catch massively on a daily basis during the reproductive season [50]. Here, we show genetic, morphometrical and demographic evidence for a substantial trend in monogamous mating in *W. scintillans*, which provides first reported case in cephalopods.

## 2. Methods

### (a) Animal and embryo collection

The squid, *W. scintillans* was obtained from fishery catches by bottom trawls towed around the Oki island (Shimane Prefecture) and Sakai-port off (Tottori Prefecture), or by fixed net set around and near the shelf break in the innermost part of the Toyama bay (Toyama Prefecture), Japan. The commercial fishing period of this species is approximately from January to May in Shimane/Tottori, and from March to May in Toyama. All the morphometric measurements were undertaken on specimens collected in Shimane/Tottori during 2016-2019, whereas for microsatellite analysis of the spermatangia and the parentage analysis, live animals caught at Toyama bay were used. At the Uozu Aquarium, spawning was induced at 14°C in dark. Spawning occurred spontaneously in the aquarium tank when the field-caught females were transported immediately after the catch. Each egg-string retrieved in isolation was then cultured for 4-5 days at 17°C until hatching and thereafter, paralarvae being picked at random (15∼45 specimens/egg-string, n=4) were genotyped.

### (b) Measurements of growth and reproductive indices

The squid specimens were measured (mostly within one day after fishing) for dorsal mantle length (ML), total wet weight (body weight; BW), testis weight (TW), spermatophoric complex weight (SCW), number of spermatophore stored in the Needham’s sac and the terminal organ, ovary weight (OW) and number of spermatangium on the left and right side of the female seminal receptacle. The number of spermatangium was counted by viewing under a stereomicroscope. Testicularsomatic index (TSI) and ovariansomatic index (OSI) were calculated as TSI = 100 x TW x BW^-1^ and OSI = 100 x OW x BW^-1^, respectively. To score the number of spermatophore, spermatophoric complex was fixed in 10% formalin in seawater, thereafter dissected under a stereoscope. All males of *W. scintillans* were mature (stage V or VI according to classification by [51]). The data for other squid species were extracted from the literature (Table S1). Average spermatophore weight (ASW) was estimated from the regression lines of Figs. 2*B* and 2*C* and in each individual, gonadal investment (resource allocation) to spermatophore was calculated as 100 x (SpN x ASW) x (TW + SCW) ^-1^, where SpN indicates number of spermatophore stored in the spermatophoric sac. Resource allocation to testis was calculated as 100 x TW x (TW + SCW)^-1^. TSI and other morphometric data of *I. paradoxus* [37], *H. bleekeri* [52] and *Doryteuthis plei* [53] were provided from the corresponding authors of previous reports.

**Fig. 1.**
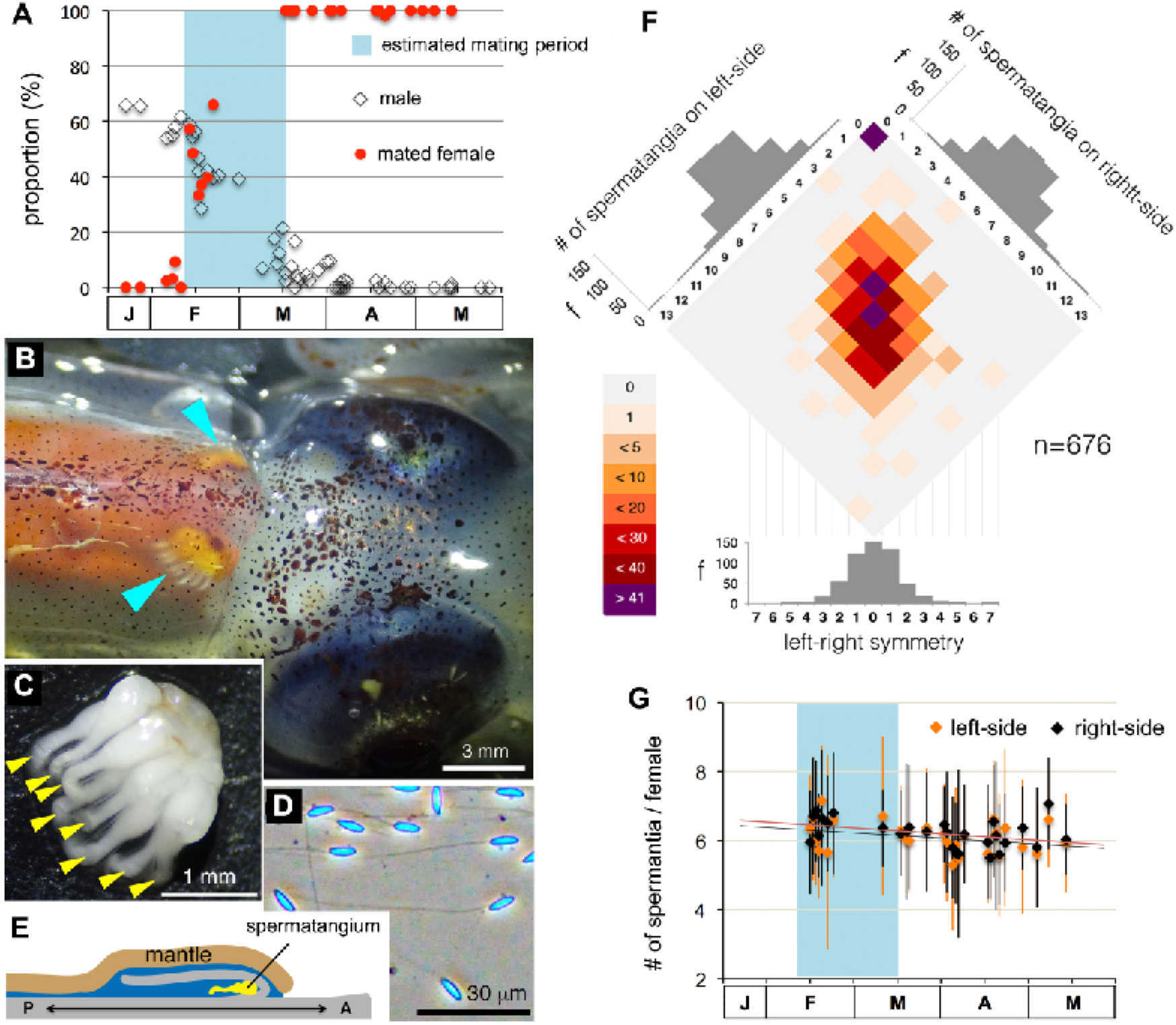
The mode of and change in sperm storage throughout the reproductive season in *W. scintillans*. (*A)* Population dynamics of males and mated females from January to May. A *cyan* box indicates the estimated mating period. (*B*) A dorsal view of the female around the neck. *Arrowheads* point to the spermatangia visible through the transparent mantle. (*C*) A single mass of spermatangium with unidirectionally oriented ejaculatory ducts (arrowheads). (*D*) Spermatozoa stored in the spermatangia. (*E*) An anatomical illustration of the female seminal receptacle along the body axis (A, anterior; P, posterior). (*F*) The frequency of female individuals having different numbers of spermatangia in bilateral storage sites. (*G*) A seasonal change in spermatangia number in females.

**Fig. 2.**
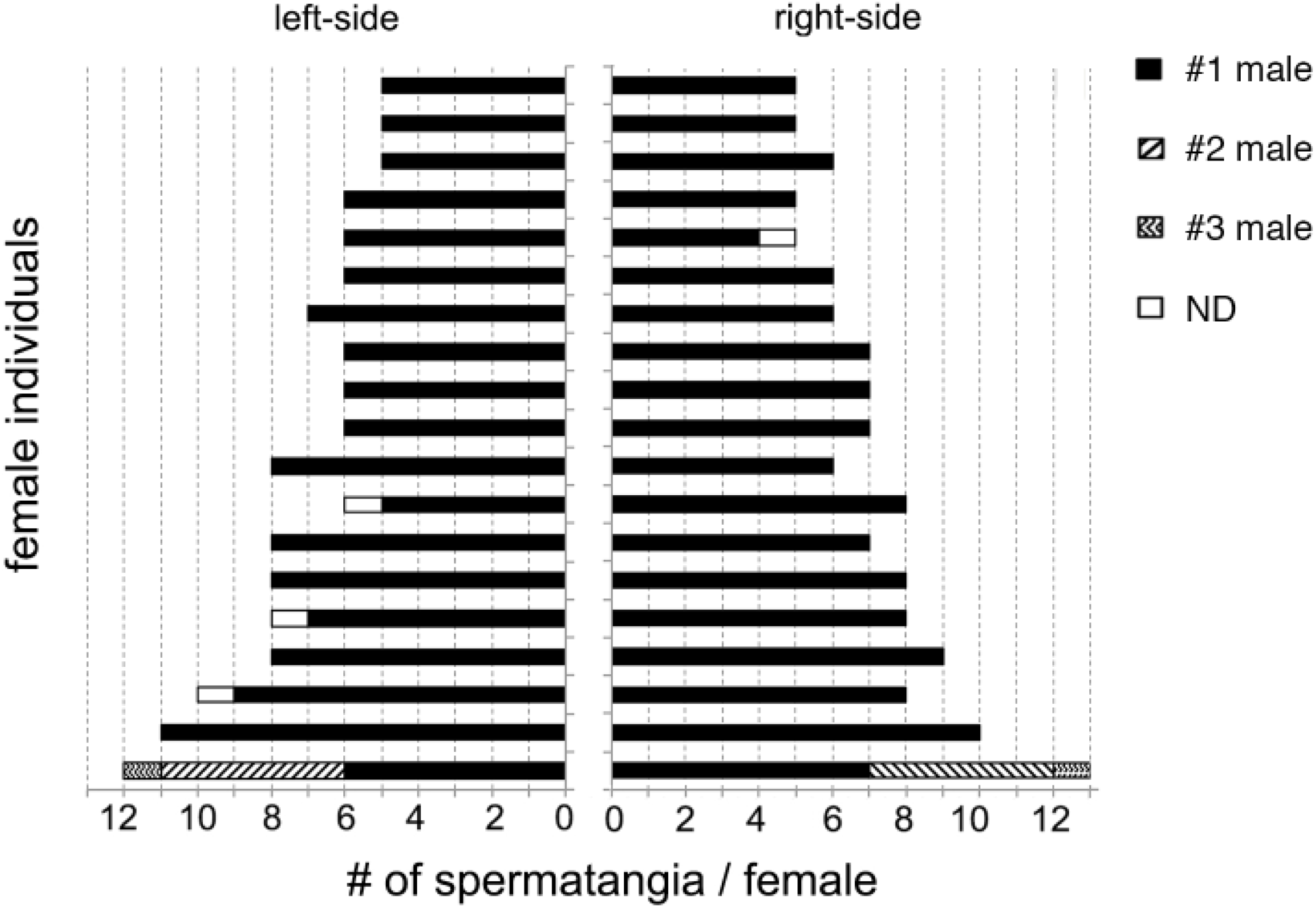
Paternity analysis of each spermatangium stored on the females. Using a total of nineteen females, every single spermatangium was isolated and genotyped. Each color represents paternity share by the first (*blue*), second (*green*) and third (*red*) male. *Cyan* indicates unidentified paternity.

### (c) Development of microsatellite markers

Microsatellite makers (GenBank Acc. LC514114 - LC514117) are developed from partial genome sequences of *W. scintillans* obtained by next-generation sequencing shared by National Center for Child Health and Development in Tokyo. The MISA pipeline [54] was used to identify reads containing microsatellite repeats and primer sets to amplify the repeats. We searched for trimers, tetramers, pentamers and hexamers with sufficient flanking sequences to develop PCR primers. A total of 1632 putative SSRs with repeat size > 60bp and flanking regions at the both ends were detected and for SSR screening, thereafter we chose 20 primer pairs flanking tetra- and trinucleotide SSR motif with a minimum of 20 repetitions and the expected product size between 140 and 280 bp. Primer3 [55] was implemented to predict primers pairs targeting the flanking regions. PCR amplification was carried out with genomic DNAs isolated from five different individuals and validated by agarose gel electrophoresis. The primer pairs that gave no band or no obvious polymorphism in band size were eliminated. The remains were further analyzed by acrylamide gel electrophoresis with a dozen of DNA samples obtained from different individuals. Consequently, we selected four SSR loci that were fully characterized by fragment length analysis (see below). Nucleotide sequences of microsatellite makers and their characteristics are shown in Table S2.

### (d) Paternity analysis

A mass of spermatangia was removed from the female seminal receptacle, soaked in 70% ethanol for 10 min and disassembled into single spermatandium with fine forceps. Each spermatangium was placed in the 1.5ml test tube, and digested with 50 μl of lysis buffer (10 mM Tris-Cl (pH 8), 500mM NaCl, 50mM EDTA, 2% dithiothreitol, 1 mg/ml Proteinase K (Sigma-Aldrich)) at 52° °C for overnight with constant agitation (120 rpm). After centrifugation (14,000 rpm, 10min, 4°C), the supernatant was transferred into new test tube followed by phenol/chloroform extraction. Extracted DNA was precipitated by adding 1/10 volume of 3 M sodium acetate (pH 5.2) and x2.5 volume of 99.5% ethanol, and centrifuged (14,000 rpm, 10 min, 4°C). After removing the supernatant, the precipitant was washed with 70% ethanol, air-dried, dissolved in 50 μl of water and stored at -20°C. Isolation of genomic DNAs from mantle tissue, hatched paralavae were carried out as well. A parentage DNA analysis was carried out by genotyping the mother, her storing spermatangia, and spawned eggs, using the software program COLONY v.2.0.6.5 [56].

All genomic DNA samples were run on 1% agarose gel electrophoresis for evaluating quality and quantity. It should be noted that in rare but certain occasion, no DNA was recovered from the spermatangium, which could be explained by the absence of spermatozoa in the spermatangium. This lack of complete recovery of genomic DNA was reflected on the result that the percent of success in PCR amplification with microsatellite markers was 99.0% (1172/1184) and overall success of paternity identification was 98.9% (270/273).

Amplification of microsatellite loci was performed with polymerase chain reaction with fluorescently tagged forward primer and non-labeled reverse primer, and approximately 20 ng of genome DNA in 10 μl of reaction mixture (TaKaRa Ex Taq system) with heat denaturing (95°C, 1min) followed by 30 cycles of denaturing (9°C, 30 sec), annealing (56°C, 30 sec) and extension (72°C, 20 sec) and terminated by 15 min incubation at 72°C. Each PCR product was run on 8% mini-slab polyacrylamide gel electrophoresis (Bio-Rad, 130 V constant, 60 min), and gels stained with 0.1 μg/ml ethidium bromide and image processed by ImageQuant LAS 500 (GE Healthcare). After validation of quality, four PCR products (labeled with FAM, Hex, Cy3 and PET) amplified with the same genome DNA were mixed and subjected to fragment length analysis (3130xl Genetic Analyzer, 500LIZ, FASMAC Co. Japan). Then, by using Cervus 3.0 ([57], the combined non-exclusion probabilities for all microsatellite loci were calculated to be < 0.0001, which is sufficiently low value for correctly identifying the real sire [58].

## 3. Results

### (a) Seasonal dynamics of population and reproduction in *W. scintillans*

Females of the firefly squid, *W. scintillans* come to the shallow water to spawn in the spring. While males disappear much earlier from the coastal zones than do females [50], which is accounted for sex difference in the lifespan being approximately one month shorter in males than females [49, 59]. The coincidental disappearance of males and maiden females, occurring from mid-February to mid-March (hereafter designated as the estimated mating period, EMP), supports this scenario as being plausible (Fig. 1A). Hence, females must internally preserve spermatozoa for a considerable time until spawning. Sperm storage occurs in the male-derived spermatangia, which are affixed to the female seminal receptacle located under the collar on the bilateral sides of the nuchal cartilage (Figs. 1B–1E). Noteworthy, an equal number of spermatangia, approximately six (left side, 5.89 ± 1.58; right side, 5.96 ± 1.65; paired t test, t = –1.02, df = 582, p = 0.31), was stored on each side of the seminal receptacle (Fig. 1F). This pattern remained unaltered with a constant and gradual decrease in the number of spermatangia throughout the reproductive season (days required to lose one spermatangium from either the left- or right side was 175.4 or 192.3, respectively; Fig. 1G), suggesting a lifelong preservation of the spermatangia once attached to the female seminal receptacle.

### (b) Evidence for behavioural and genetic monogamy

This well-organized pattern of sperm storage with a left–right symmetry and an approximately fixed number led us to test the hypothesis that females receive spermatozoa by a single copulation attempt from the male (i.e., behavioral monogamy). By genotyping with validated microsatellite markers (Table S2), we found that in 94.7% (18/19) of females, all the spermatangia stored were delivered by a single male. In one case, where the number of spermatangia stored was extraordinarily large (left, 13; right, 12), three males were involved in copulation (Fig. 2).

To ascertain whether females use spermatozoa from the storing spermatangia to fertilize their eggs (i.e., genetic monogamy), a parentage DNA analysis was carried out with SSR loci of the mother, her storing spermatangia, and her brood. The COLONY analysis [56] indicated that all paralarvae analyzed were assigned to the single paternity derived from the storing spermatangia (Datasheet S1, combined non-exclusion probability < 0.001; paternity rate by a candidate male = 100%, n = 4). These results also rule out the possibility that females commit cryptic sperm storage following extra-pair copulations.

### (c) Estimation of male mating opportunities

We next performed morphometric and quantitative measurements of maturity and fecundity of *W. scintillans* individuals before, during, and after EMP. We found that males continue to accumulate spermatophores in their storage organ (Needham’s sac) throughout the season except during EMP (Fig. 3A), when the spermatophores are being used. Such a continuous increase in stored spermatophore can be explained by a higher manufacturing rate than their consumption rate, suggesting that males would have lost their copulation opportunities after EMP. The mean number of male-stored spermatophores just before EMP (pre-EMP) was ∼30, which estimates that males can copulate no greater than 2∼3 times (because the mean number of female-received spermatophores is 12). At pre-EMP, males become fully mature with the highest testicularsomatic index (TSI), an indicator of sperm-producing capacity or promiscuity (Figs. 3B–D), whereas females have just begun maturing and require several more weeks to become more fecund (Fig. 2E). However, the TSI was, to our current knowledge, extremely lower in *W. scintillans* (0.15 ± 0.09, n = 408) than in other investigated cephalopods (Fig. 3F). From another point of view, *W. scintillans* males invest most gonadal expenditure to production of the spermatangia rather than sperm, giving a stark contrast to the case of *I. paradoxus* (Figs. 3G–L) that copulates multiple times throughout the reproductive season [37]. Collectively, we assume that males of *W. scintillans* are incapable of attempting multiple mates, hence resulting in monogynous mating pattern, because of low sperm production capacity and limited mating opportunities due to male-biased operational sex ratio and absence of female remating.

**Fig. 3.**
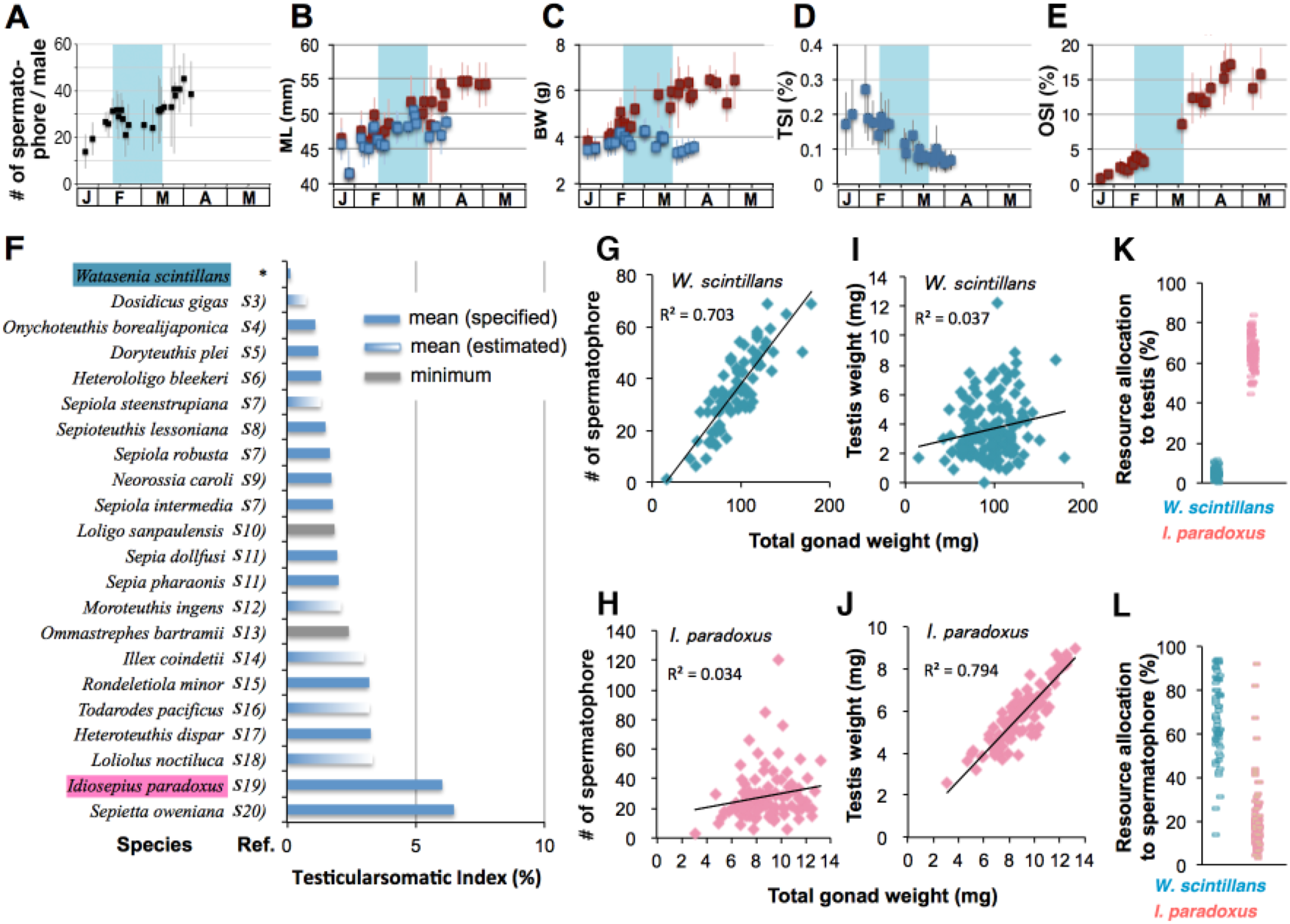
Allocation of male reproductive resources in *W. scintillans* and other squids. (*A-E*) In *W. scintillans*, seasonal changes in the number of male-storing spermatophore (*A*), mantle length (*B*), body weight (*C*), TSI (*D*) and OSI (*E*) are plotted as the mean ± SE. Data points represent males in *blue* and females in *red. Cyan* boxes indicate the estimated mating period. (*F*) A graph showing a comparison of male testicularsomatic indices (TSI) among previously reported cephalopod species and *W. scintillans* found in this study (*). Data were extracted from the literature indicated by reference # (Table S1). The columns indicate the mean (*blue column*) or minimum (*gray column*) values specified in the literature, otherwise mean values were estimated (*gradient column*) from presented data points in the graphs. (*G-H*) The GSI values from male individuals are plotted against number of spermatophore (*G, H*) or testis weight (*I, J*) in *W. scintillans* (*G, I*) and highly promiscuous *Idiosepius paradoxus* (*H, J*). (*K, L*) In each individual, allocation of male reproductive resources to testis (*K*) or spermatophore (*L*) is expressed in percentage.

## 4. Discussion

One of the hallmarks of reproduction in most coleoid cephalopods is their copulation behavior by which males deposit sperm packages (spermatophores) on female using the heterocotylated arm(s). Because females are primarily promiscuous, the spermatangia are delivered simultaneously or sequentially from multiple males, resulting in random distribution of mixed spermatangia around the deposition site on the female. However, we contingently found that females of *W. scintillans* store masses of the spermatangia that are evenly distributed at exact locations on bilateral sides of the nuchal cartilage with approximately six on each side. Such the extraordinary regular pattern of spermatangium placement was unusual in cephalopods, thus prompted us to test whether all these spermatangia were derived from a single male.

Our current genetic data (paternity of each spermatangium stored on a female) clearly demonstrated that most females of squid *W. scintillans* mate with only one male (Fig. 2), therefore suggesting behavioral monoandry (females mate with only one partner). Furthermore, parentage analysis identified no DNA mismatch between the spermatangia and embryos from the same females, confirming genetic monogamy (Fig. S1). During reproductive season, females may spawn eggs several times at certain intervals [49], however other males would not be involved in replenishing the spermatangia during the intervals because 1) males disappear before females being fully fecund (Figs. 1A, 3E), and 2) spontaneous loss of once-attached spermatangium to female seldom occurs (Fig. 1G). Accordingly, male mating opportunities would be limited due to infrequent female remating and male-biased sex ratio. Both tertiary (adult) sex ratio and operational sex ratio [60] were largely male-biased in the beginning of and throughout EMP, respectively (Fig. 1A). Under these conditions, mathematical modeling predicts that monogyny (males mate with only one partner) can become fixed as evolutionary stable strategy (ESS)[60, 61]. So, we speculate that female monoandry had first established followed by male monogyny, which together brought into mutual monogamy [60]. If male monogyny becomes ESS, then evolution could favor males who invest more energy on something other than fecundity (sperm production) [62, 63], which explains the observed least investments on testis (sperm production) and spermatangium (copulation opportunity) in this species. Instead of allocating their reproductive investment toward greater fecundity, males would have chosen a “live-fast-die-young” life-history strategy to adapt to a mechanism of first-male sperm precedence [64] (Fig. 1A). We speculate that the low level of male fecundity had occurred as the consequence of female-driven monoandry [62, 63].

If females and males are predominantly pursuing a monogamous mating strategy, what could be the benefit they reap in exchange for eroding the genetic diversity of the offspring? And in doing so, how do they physiologically evade remating attempts despite physically being together? Therefore, the first question arises as to what was a feasible cause or causes that drove their mating system to be primarily, if not exclusively, monoandrous. It is unambiguous that some factors known to involve monogamy in other taxa do not work for this species such as long-term pair-bond, biparental care, low population density, habitat limitation, time constraint for reproduction and enforcement (mate guarding) [16, 18, 21, 65], simply because they make condensed spawning migrations and males disappear entirely before peaking at egg laying. Alternatively, considering that this species suffers from high predation pressures and serves as a dominant pray for demersal fishes [66], a time-consuming courtship behavior could potentially be deleterious to their survival; hence, multiple mating would reduce reproductive success for both sexes [8, 67, 68]. There are ample examples to support the prediction that a predation risk serves an evolutionally force to monoandry [8]. For example, in the cicada *Subpsaltria yangi*, when males emit an advertising call, only virgin females make a response call, thereafter males fly to them to mate (hence monoandry). These flights by males are at high risk – only 25% of the second flights were successful and the remaining failures were mostly attacked by the robber fly, *Philonicus albiceps* [69].

It is widespread phenomena across taxa that non-virgin females lose sexual attractiveness or responsiveness to male-specific courtship signaling, and the underlying mechanisms may vary with physiological, physical and behavioral changes upon mating [70-72]. Nevertheless, their fitness consequences would be mostly against the risk of predation. Although the actual mating behaviors of deep-sea organisms are largely concealed, mating of any species should be well adopted in the dark environment [48]. The communication-like bioluminescence signaling executed by the Octopodiformes squid, *Taningia danae* was regarded as a potential courtship behavior [73]. *W. scintillans* has special eyes (photoreceptor cells) containing three visual pigments (λ_max_ ∼471, ∼484 and ∼500 nm), possibly allowing them to distinguish conspecific illumination (green) from environmental down-welling light (blue) [74]. It is therefore feasible that bioluminescence of the firefly squid could play a role for courtship signaling or mate search in a once-in-a-lifetime fashion. In this study, the fine-scale analysis of seasonal dynamics in demographics, mating status and reproductive indices of each individual from fishery catches sculptured the reproductive landscape of this species. In summary, we provide, for the first time in cephalopods, genetic evidence for virtually monoandrous mating pattern and its adaptive significance awaits deciphering in the light of ecological niche of this species.

## Data accessibility

All datasets for information of the SSR markers and the paternity analysis are available in the electronic supplementary material files and newly developed SSR markers have been deposited at the DDBJ/GenBank with accession no. LC514114 - LC514117.

## Authors’ contributions

NH conceived this study. NS, SIT, NEA, SK, TS and NH performed experiments. NS, YI, OI, MAY and NH analyzed data. NH wrote the paper.

## Acknowledgments

We thank Mitsuhiro Fuwa, Tomoharu Kimura, Mari Saito, Kazusa Saiba and other members of Uozu aquarium for helping sample collection, Koichi Ozaki for the lab facility. We thank Lígia H. Apostólico and José E. A. R. Marian (Univ. of Sao Paulo) for providing individual data for *D. plei*.

## Supplemental data

**Table S1.** Reference for testicularsomatic index of various squid species used in Fig. 3F.

| ref# |  |
| --- | --- |
| S3 | Markaida U. and Sosa-Nishizaki O. (2001) Reproductive biology of jumbo squid *Dosidicus gigas* in the Gulf of California, 1995–1997. Fish. Res. 54, 63-82. |
| S4 | Murata M., Ishii M. and Osako S. Some Information on Copulation of the Oceanic Squid *Onychoteuthis borealijaponica* Okada. (1982) Bull. Jap. Soc. Sci. Fish. 48,351-354. |
| S5 | Apostólico L.H. and Marian J.E.A.R. (2018) Dimorphic male squid show differential gonadal and ejaculate expenditure. Hydrobiologia 808, 5-22. |
| S6 | Iwata Y., Munehara H., Sakurai Y. (2005) Dependence of paternity rates on alternative reproductive behaviors in the squid *Loligo bleekeri*, Mar. Ecol. Prog. Ser. 298, 219-228. |
| S7 | Salman A. and Onsoy, B. (2004) Analysis of fecundity of some bobtail squid of the genus *Sepiola* (Cephalopoda: Sepiolida) in the Aegean Sea (eastern Mediterranean). J Mar. Biol. Ass. UK 84, 781-782. |
| S8 | Thulasitha W.S., Charles G.A., and Sivashanthini K. (2010) Reproductive Characteristics of Squid *Sepioteuthis lessoniana* (Lesson, 1830) from the Northern Coast of Sri Lanka. J. Fish. Aqu. Sci. 5, 12-22. |
| S9 | Cuccu D., Mereu M., Masala P., Cau A. and Jere P. (2011) Male reproductive system in *Neorossia caroli* (Joubin 1902) (Cephalopoda: Sepiolidae) from Sardinian waters (western Mediterranean Sea) with particular reference to sexual products. Invert. Reprod. Develop. 55, 16-21. |
| S10 | Costa P.A.S., Fernandes F.C. (1993) Reproductive cycle of *Loligo sanpa ulensis* (Cephalopoda: Loliginidae) in the Cabo Frio region, Brazil. Mar. Eco. Pro. Ser. 101, 91-97. |
| S11 | Gabr H.R., Hanlon R.T., Hanafy M.H. and El-Etreby S.G. (1998) Maturation, fecundity and seasonality of reproduction of two commercially valuable cuttlefish, *Sepia pharaonis* and *Sepia dollfusi* in the Suez Canal. Fish. Res. 36, 99-115. |
| S12 | Jackson G.D., Semmens J.M., Phillips K. L., Jackson C.H. (2004) Reproduction in the deepwater squid *Moroteuthis ingens*, what does it cost? Mar. Biol. 145, 905–916. |
| S13 | MARCELAL B., ANÍBALAUBONE I. and PASCUAL L. (2006) REPRODUCTIVE BIOLOGY OF RED SQUID (*Ommastrephes bartramii*) IN THE SOUTHWEST ATLANTIC NORMAE. Rev. Invest. Desarr. Pesq. N° 18, 5-19. |
| S14 | Cerola L. and Milone N. (2017) Growth and reproduction of the squid *Illex coindetii* Verany, 1839 in the southern Adriatic Sea, central Mediterranean. Medit. Mar. Sci., 18, 107-120. |
| S15 | Onsoy B., Ceylan B. and Salman A. (2013) Reproductive biology of the lentil bobtail squid, *Rondeletiola minor* (Cephalopoda: Sepiolidae) from the eastern Mediterranean. 93, 851-854. |
| S16 | Ikeda Y., Sakurai Y. and Shimazaki K. (1991) Development of Male Reproductive Organs During Sexual Maturation in the Japanese Common Squid *Todarodes pacificus*. Nippon Suisan Gakkaishi 57, 2237-2241. |
| S17 | Hoving H.J.T. and Laptikhovsky V. (2008) Reproduction in *Heteroteuthis dispar* (Rüppell, 1844) (Mollusca: Cephalopoda): A sepiolid reproductive adaptation to an oceanic lifestyle. Mar. Biol. 154, 219-230. |
| S18 | Jackson G.D. and Moltschanivskyj N.A. (2001) Temporal variation in growth rates and reproductive parameters in the small near-shore tropical squid *Loliolus noctiluca*; is cooler better? Mar. Ecol. Prog. Ser. 218,167-177. |
| S19 | Sato N., Yoshida MA. and Kasugai T. (2017) Impact of cryptic female choice on insemination success: Larger sized and longer copulating male squid ejaculate more, but females influence insemination success by removing spermatangia. Evolution 71, 111-120. |
| S20 | Giordano D. et al. (2009) Distribution and biology of *Sepietta oweniana* (Pfeffer, 1908) (Cephalopoda: Sepiolidae) in the Southern Tyrrhenian Sea (Central Mediterranean Sea), Cah. Biol. Mar. 50, 1-10. |

**Table S2.**
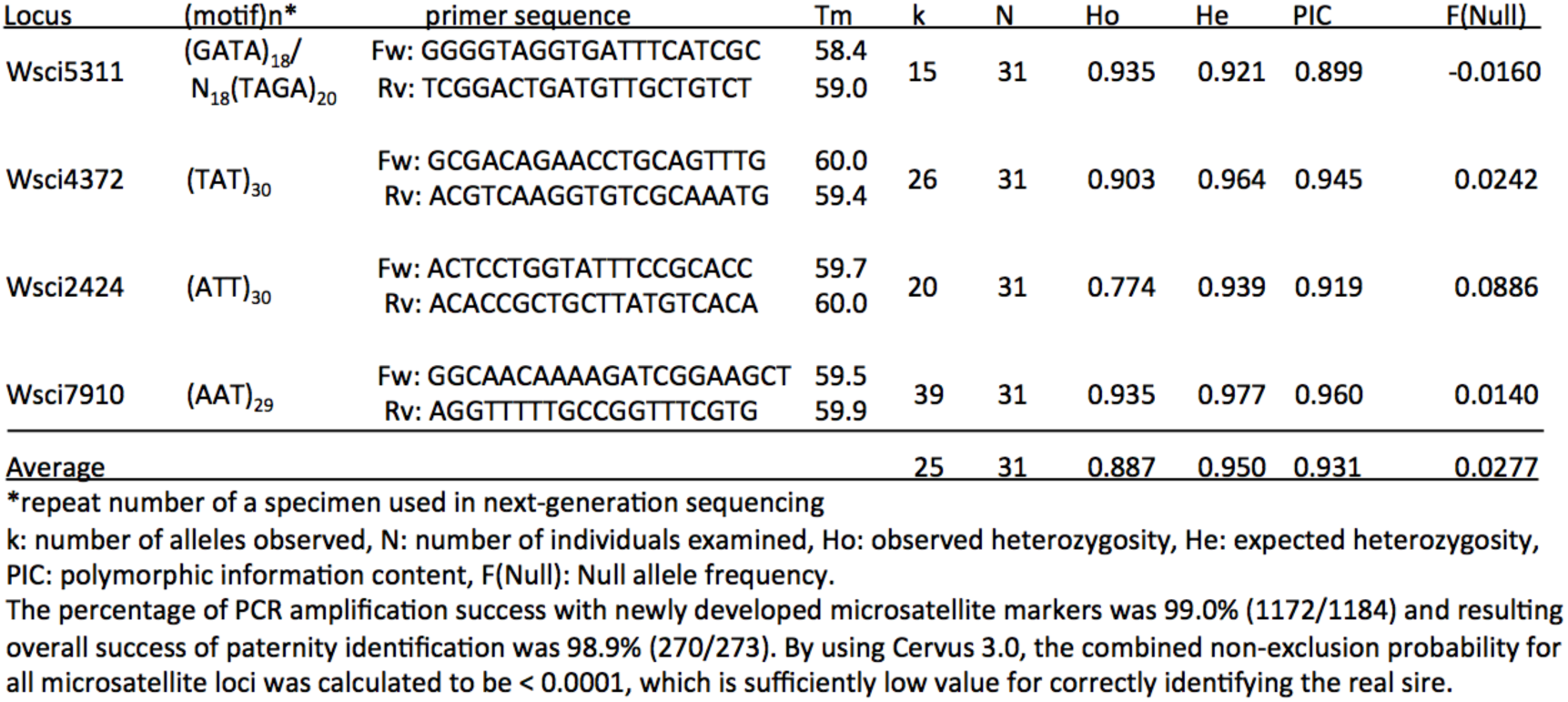
Information of microsatellite makers for *Wastasenia scintillans*.

## Datasheet S1

| Experiment 1 | Wsci2424 |  | Wsci4372 |  | Wsci7910 |  |
| --- | --- | --- | --- | --- | --- | --- |
| female mantle | 211 | 215 | 135 | 135 | 204 | 216 |
| sperm mass | 197 | 208 | 135 | 156 | 114 | 180 |
| embryo #2 | 197 | 215 | 135 | 156 | 114 | 216 |
| embryo #3 | 208 | 215 | 135 | 135 | 180 | 216 |
| embryo #4 | 197 | 215 | 135 | 135 | 114 | 216 |
| embryo #5 | 208 | 212 | 135 | 135 | 180 | 204 |
| embryo #6 | 197 | 215 | 135 | 156 | 114 | 216 |
| embryo #7 | 197 | 215 | 135 | 156 | 114 | 216 |
| embryo #8 | 197 | 212 | 135 | 135 | 114 | 204 |
| embryo #9 | 208 | 212 | 135 | 156 | 180 | 204 |
| embryo #11 | 208 | 212 | 135 | 156 | 180 | 204 |
| embryo #13 | 197 | 212 | 135 | 156 | 114 | 204 |
| embryo #15 | 197 | 215 | 135 | 156 | 114 | 216 |
| embryo #16 | 208 | 212 | 135 | 156 | 180 | 204 |
| embryo #17 | 208 | 215 | 135 | 135 | 180 | 216 |
| embryo #18 | 197 | 212 | 135 | 156 | 114 | 204 |
| embryo #19 | 197 | 215 | 135 | 156 | 114 | 216 |

| Experiment 2 | Wsci2424 |  | Wsci7910 |  |
| --- | --- | --- | --- | --- |
| female mantle | 173 | 185 | 171 | 318 |
| sperm mass | 191 | 216 | 105 | 180 |
| embryo #1 | 185 | 215 | 105 | 318 |
| embryo #2 | 173 | 215 | 105 | 171 |
| embryo #3 | 185 | 215 | 105 | 318 |
| embryo #4 | 185 | 191 | 180 | 318 |
| embryo #5 | 185 | 191 | 180 | 318 |
| embryo #7 | 185 | 191 | 180 | 318 |
| embryo #8 | 173 | 215 | 105 | 171 |
| embryo #9 | 173 | 215 | 105 | 171 |
| embryo #10 | 185 | 191 | 180 | 318 |
| embryo #11 | 185 | 215 | 105 | 171 |
| embryo #12 | 173 | 215 | 105 | 171 |
| embryo #13 | 173 | 215 | 105 | 171 |
| embryo #14 | 173 | 191 | 171 | 180 |
| embryo #15 | 173 | 191 | 171 | 180 |
| embryo #16 | 185 | 191 | 180 | 318 |
| embryo #17 | 185 | 215 | 180 | 318 |
| embryo #18 | 185 | 215 | 105 | 318 |
| embryo #19 | 185 | 191 | 180 | 318 |
| embryo #20 | 185 | 191 | 180 | 318 |

| Experiment 3 | Wsci2424 |  | Wsci5311 |  | Wsci4372 |  |
| --- | --- | --- | --- | --- | --- | --- |
| female mantle | 200 | 204 | 146 | 208 | 116 | 166 |
| sperm mass | 182 | 200 | 170 | 203 | 107 | 123 |
| embryo #1 | 182 | 200 | 170 | 208 | 107 | 116 |
| embryo #3 | 200 | 204 | 170 | 208 | 122 | 166 |
| embryo #4 | 200 | 200 | 146 | 203 | 107 | 116 |
| embryo #5 | 200 | 204 | 146 | 170 | 107 | 116 |
| embryo #7 | 200 | 204 | 170 | 208 | 107 | 116 |
| embryo #8 | 182 | 204 | 146 | 203 | 122 | 166 |
| embryo #9 | 182 | 204 | 203 | 207 | 107 | 116 |
| embryo #10 | 200 | 204 | 170 | 208 | 107 | 116 |
| embryo #11 | 182 | 200 | 146 | 203 | 107 | 166 |
| embryo #12 | 200 | 200 | 146 | 170 | 107 | 166 |
| embryo #13 | 200 | 200 | 203 | 208 | 116 | 122 |
| embryo #14 | 182 | 200 | 146 | 203 | 107 | 116 |
| embryo #15 | 182 | 204 | 203 | 208 | 122 | 166 |
| embryo #16 | 182 | 204 | 170 | 208 | 166 | 122 |
| embryo #17 | 200 | 204 | 203 | 208 | 116 | 122 |
| embryo #18 | 200 | 200 | 203 | 208 | 107 | 116 |
| embryo #19 | 182 | 204 | 146 | 170 | 107 | 166 |
| embryo #20 | 200 | 204 | 203 | 208 | 116 | 122 |

| Experiment 4 | Wsci2424 |  | Wsci5311 |  | Wsci4372 |  | Wsci7910 |  |
| --- | --- | --- | --- | --- | --- | --- | --- | --- |
| female mantle | 150 | 267 | 221 | 225 | 203 | 241 | 266 | 280 |
| sperm mass | 165 | 279 | 204 | 241 | 225 | 251 | 272 | 298 |
| embryo #2 | 150 | 165 | 225 | 242 | 225 | 241 | 280 | 298 |
| embryo #3 | 267 | 279 | 204 | 225 | 203 | 251 | 266 | 272 |
| embryo #4 | 165 | 267 | 221 | 242 | 225 | 241 | 266 | 272 |
| embryo #5 | 150 | 279 | 204 | 225 | 203 | 225 | 272 | 280 |
| embryo #6 | 150 | 279 | 205 | 225 | 203 | 225 | 272 | 280 |
| embryo #7 | 150 | 279 | 205 | 225 | 241 | 250 | 272 | 280 |
| embryo #8 | 150 | 165 | 205 | 221 | 203 | 251 | 280 | 298 |
| embryo #9 | 150 | 165 | 204 | 221 | 203 | 251 | 280 | 298 |
| embryo #10 | 165 | 267 | 221 | 242 | 225 | 241 | 266 | 298 |
| embryo #11 | 150 | 279 | 221 | 241 | 241 | 251 | 272 | 280 |
| embryo #12 | 150 | 279 | 225 | 242 | 203 | 251 | 272 | 280 |
| embryo #13 | 150 | 279 | 225 | 241 | 203 | 225 | 272 | 280 |
| embryo #14 | 267 | 279 | 205 | 225 | 203 | 251 | 272 | 280 |
| embryo #15 | 150 | 279 | 221 | 242 | 241 | 251 | 272 | 280 |
| embryo #16 | 150 | 165 | 204 | 225 | 203 | 225 | 280 | 298 |
| embryo #17 | 165 | 267 | 221 | 242 | 203 | 225 | 266 | 298 |
| embryo #18 | 165 | 267 | 221 | 242 | 225 | 241 | 266 | 298 |
| embryo #19 | 267 | 279 | 221 | 242 | 203 | 225 | 272 | 280 |
| embryo #20 | 150 | 279 | 204 | 225 | 203 | 251 | 272 | 280 |
| embryo #21 | 165 | 267 | 221 | 241 | 225 | 241 | 280 | 298 |
| embryo #22 | 267 | 279 | 221 | 242 | 225 | 241 | 266 | 272 |
| embryo #23 | 150 | 165 | 221 | 242 | 203 | 251 | 280 | 298 |
| embryo #24 | 150 | 279 | 225 | 242 | 241 | 251 | 272 | 280 |
| embryo #25 | 267 | 278 | 205 | 221 | 204 | 251 | 266 | 272 |
| embryo #26 | 165 | 267 | 225 | 242 | 204 | 251 | 266 | 298 |
| embryo #27 | 150 | 279 | 221 | 242 | 225 | 241 | 272 | 280 |
| embryo #28 | 150 | 279 | 225 | 241 | 241 | 251 | 280 | 298 |
| embryo #29 | 165 | 267 | 204 | 221 | 225 | 241 | 266 | 298 |
| embryo #30 | 150 | 165 | 204 | 221 | 241 | 251 | 280 | 298 |
| embryo #31 | 150 | 279 | 204 | 225 | 225 | 241 | 272 | 280 |
| embryo #32 | 267 | 279 | 204 | 221 | 241 | 251 | 266 | 272 |
| embryo #33 | 150 | 165 | 221 | 242 | 241 | 251 | 280 | 298 |
| embryo #34 | 165 | 267 | 204 | 225 | 225 | 241 | 266 | 298 |
| embryo #35 | 165 | 268 | 204 | 225 | 241 | 251 | 266 | 298 |
| embryo #37 | 150 | 279 | 221 | 242 | 241 | 251 | 272 | 280 |
| embryo #38 | 165 | 267 | 221 | 242 | 203 | 251 | 266 | 298 |
| embryo #39 | 165 | 267 | 221 | 242 | 203 | 251 | 280 | 298 |
| embryo #40 | 165 | 267 | 225 | 242 | 203 | 222 | 266 | 298 |
| embryo #41 | 267 | 279 | 204 | 225 | 225 | 241 | 266 | 272 |
| embryo #43 | 150 | 165 | 225 | 242 | 225 | 241 | 280 | 298 |
| embryo #44 | 150 | 279 | 204 | 221 | 241 | 251 | 272 | 280 |
| embryo #45 | 267 | 279 | 204 | 221 | 203 | 225 | 266 | 272 |
| embryo #46 | 150 | 165 | 225 | 241 | 225 | 241 | 280 | 298 |
| embryo #47 | 267 | 279 | 204 | 221 | 203 | 225 | 266 | 298 |
| embryo #48 | 150 | 279 | 204 | 221 | 203 | 225 | 272 | 280 |

## Datasheet S1 Parentage analysis

Genotyping was carried out with the mother, her stored spermatangia and spawned eggs using four SSR markers (n = 4). All hatched paralarvae, every single spermatangium, i.e., six on left side, six on right side, stored by the female and her mantle tissue were subjected to DNA extraction followed by PCR and then fragment analysis. Microsatellite null alleles were eliminated from the analysis. The estimated size (bp) of each amplicon was shown. The data clearly show that in all microsatellite loci amplified from paralarval samples, one of two alleles was derived from spermatozoa stored in the spermatangia, and the other from the mother.

